# Lipid-mediated organization of prestin in the outer hair cell membrane and its implications in sound amplification

**DOI:** 10.1101/2022.05.30.494028

**Authors:** Sepehr Dehghani-Ghahnaviyeh, Zhiyu Zhao, Emad Tajkhorshid

## Abstract

Prestin is a high-density motor protein in the outer hair cells (OHCs), whose conformational response to acoustic signals alters the shape of the cell, thereby playing a major role in sound amplification by the cochlea. Despite recent structural determination in active and inhibited states, the details of prestin’s intimate interactions with the membrane, which are central to its function remained unresolved. Here, employing a large set (collectively, more than 0.5 ms) of coarse-grained molecule dynamics simulations, we characterize the nature of prestin’s lipid-protein interactions, demonstrating their impact on the organization of prestin at densities relevant to the OHCs and its effectiveness in reshaping OHCs. Beyond local enrichment/depletion of various lipid types, prestin causes drastic anisotropic membrane deformation, which in turn mediates a preferential membrane organization of prestin in which deformation patterns by neighboring prestin copies are aligned constructively. The reduced membrane rigidity accompanying this arrangement is hypothesized to maximize the mechanical impact of prestin on OHC reshaping during cochlear sound amplification. Prestin’s preferential arrangement is further verified by extended simulations demonstrating strong correlation between prestin neighbors in their orientations. These results demonstrate a strong case of protein-protein cooperative communication in membrane, purely mediated by their interactions with lipids.

## Introduction

Hearing is a major sensory mechanism evolved to enable higher organisms to detect auditory cues from their environment.^1,2^ Species such as humans and other mammals are equipped with a complex auditory system, composed of several organs that are involved in different steps of the hearing process. Many of the complexities engineered into the hearing system have been evolved for distinguishing between different frequencies.^1,2^ A key aspect of hearing, however, is sound amplification within a spiral-shaped organ in the inner ear known as the cochlea, without which the hearing system is dysfunctional.^3^ Cochlear amplification relies on specialized cells known as the outer hair cells (OHCs) and their unique piezoelectric character.^4,5^ Modulations in the membrane potential of the OHCs result in their mechanical response in the form of vibration (a property termed “electromotility”).^4,5^ This vibration is synchronized to the sound signal and provides mechanical amplification by feeding back into the traveling wave.^4–6^ The source of electromotility in the OHCs is a motor membrane protein, called prestin, which generates somatic forces and alters the structure of the OHCs as a result of its voltage-dependent conformational changes.^7–9^ Dysfunction of prestin or other elements in the OHCs, e.g., by excessive noise or aging, can result in a reduction of sound sensitivity or even total deafness.^10–12^

Prestin is a member of the anionic transporter superfamily, SLC26A5,^13,14^ and similar to other members of the superfamily, it binds to anions (e.g., Cl^-^).^15^ However, unlike the other members, which functions as transport proteins, prestin acts as a motor protein. Despite its discovery and biochemical characterization for decades,^8,9,16,17^ the structure of prestin was resolved only recently. The cryo-EM models of prestin revealed a homodimeric architecture with a two-fold symmetry.^18,19^ The transmembrane (TM) region of each protomer is composed of two domains, the core domain (residues 80-202 and 338-432) and the gate domain (residues 209-315 and 437-497), forming an anion binding site at their interface.^18–21^

The structure of prestin has been captured with bound Cl^-^ ions (agonist), as well as with other negatively charged ligands such as the inhibitor salicylate.^18,19^ In the agonist (Cl^-^)- or inhibitor-bound structures, the core and gate domains exhibit different relative arrangements, generating two conformational/functional states coined as contracted and expanded, with a smaller protein crosssectional area in the former.^18,19^ The transition between these two conformations is hypothesized to be the source of prestin-mediated vibration of the OHCs, and, therefore, key to sound amplification in the cochlea.^18^

Freeze-fracture electron microscopy has shown a high density of prestin on the surface of the OHCs, estimating a separation of only ~200 Å between prestin dimers in the cellular membrane.^22^ The structural changes of the OHCs in response to prestin’s activation can therefore be substantial. ^15,22^ The packing ensures that even small conformational changes of individual prestin dimers will add up to sufficiently large structural changes of the whole cell, and, therefore, to effective sound amplification.^22,23^ Given its high cellular density, it is important to also understand organization of prestin dimers within the membrane, which might in turn affect their effectiveness and even cooperativity.

As described in previous studies, the interaction between the prestin and its embedding membrane is crucial for its role in sound amplification.^24–30^ Nevertheless, and despite the availability of atomic-resolution structures,^18,19^ the details of prestin’s interactions with lipids and the membrane remain largely elusive.

Lipid-protein interactions have been the subject of numerous studies, as lipids play an active role in modulating the structure and function of membrane proteins.^30–33^ Owing to their high spatial and temporal resolutions and their complementarity to experimental results, molecular dynamics (MD) simulations have played a major role in our current understanding of lipid-protein interactions in near-native membranes.^33–39^ In order to improve sampling of lipid-protein interactions, especially in heterogeneous membranes, many MD studies have taken advantage of coarse-grained (CG) representations, e.g., using Martini, a widely used CG method for lipid structures.^40,41^ Martini CG simulations of membranes and membrane proteins have contributed significantly to the understanding of how membrane proteins modulate their lipidic environments, both at the distribution and structural levels.^42,43^

Here, we have employed an extensive set of MD simulations to characterize the structural effects of prestin on the membrane. Using different simulation designs, we also ask the question whether these effect might in turn modulate prestin’s organization in the membrane at densities representing its concentration in the OHCs. The simulations capture consistently the profound effect of prestin in deforming the membrane, in a particular pattern caused by anisotropically elevating or depressing lipids at its different flanks, an effect that extends over a long range of ten nanometers away from the protein. More importantly, multiple-prestin simulations show that the patterns of membrane deformation induced by the neighboring proteins is used to sync their orientations. This through-lipid communication between the proteins provides an energetically favorable membrane organization with a lower bending modulus, which can offer a larger cellular response, and therefore more effective in producing the mechanical role of prestin in the OHCs.

## Materials and Methods

### System setup

The prestin cryo-EM structures in contracted (Cl^-^-bound) and expanded (salicylate-bound) conformations were used as the starting models to construct initial all-atom (AA) models (Fig. S1A-D).^18^ The bound ligands and modeled lipids were removed from the cryo-EM structures. Both protein structures lack a disordered region in the intracellular domain, dividing them into two polypeptide segments (residues 13-580 and 614-724, respectively). N-terminal ammonium and C-terminal carboxy caps were added to the first and last residues in each segment, respectively, employing the Psfgen plugin of VMD.^44^ All hydrogen atoms as well as the missing side chains were also added to the structures using Psfgen.

The initial AA models were then converted to Martini CG models using the Martinize protocol as described on the Martini website (http://www.cgmartini.nl/), including an elastic network restraining pairs within a 10-Å cutoff (Fig. 1A). The protein’s secondary structure was defined based on the AA models and maintained throughout the CG simulations. The CG proteins were then embedded into a lipid bilayer composed of phosphatidylcholine (PC), phosphatidylethanolamine (PE), phosphatidylglycerol (PG), phosphatidylserine (PS), phosphatidylinositol (PI), sphingomyelin (SM), and cholesterol (CHOL), at a molar ratio of 27:13:4:17:10:12:17. The initial orientation of prestin in the membrane was obtained from the OPM (Orientations of Proteins in Membranes) database.^45^

**Figure 1:**
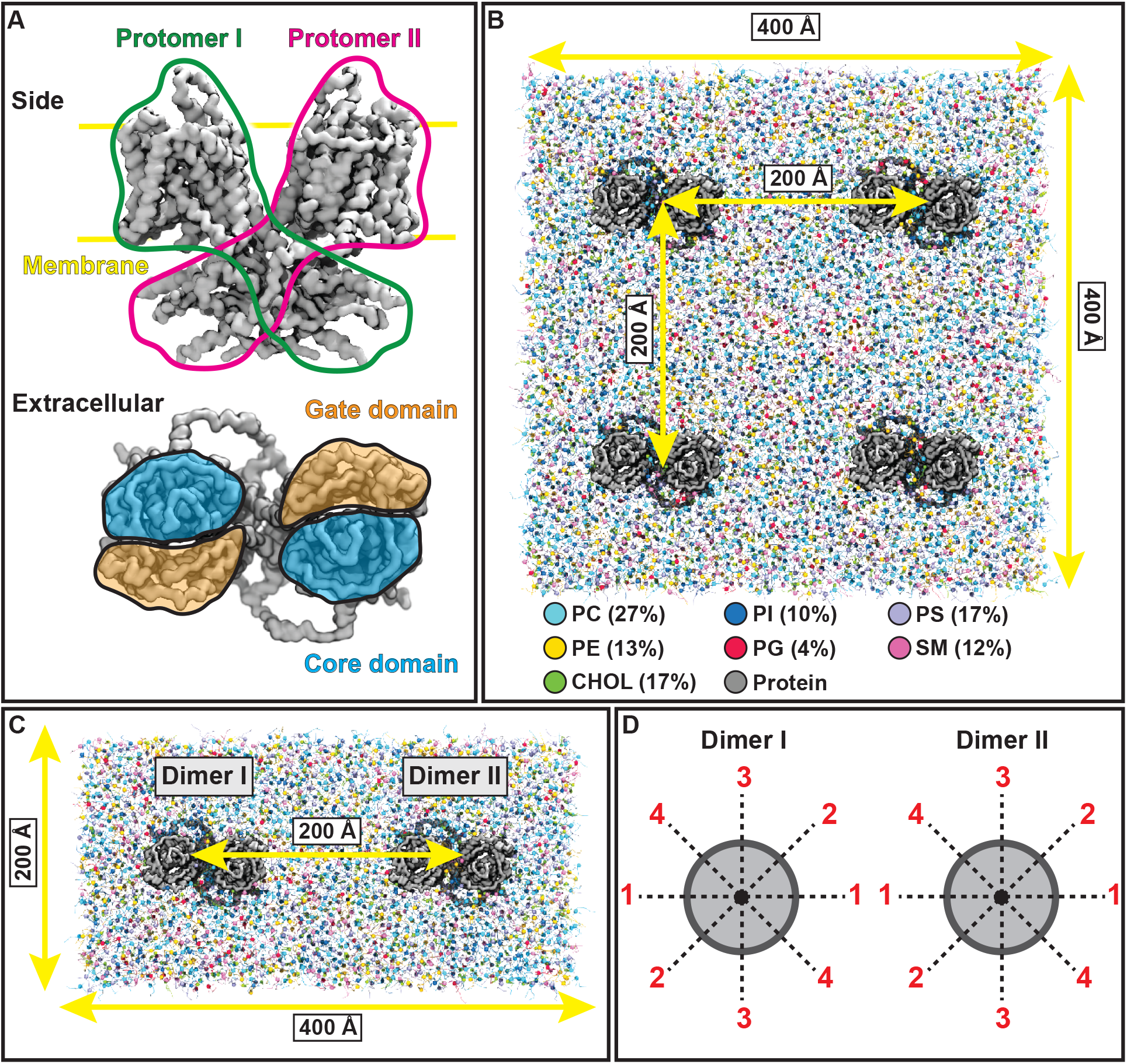
Simulation systems. **(A)** Martini-based CG model of prestin from side (top) and extracellular (bottom) views. In the top panel, the two protomers are outlined in green and magenta, respectively, and the approximate position of the membrane is marked with yellow lines. In the bottom panel, the core and gate domains are colored in orange and blue, respectively. **(B)** Simulation system to probe lipid-protein interactions, including four prestin dimers, separated by 200 Å, and embedded in a square membrane patch of side length 400 Å. The composition of the membrane is shown in the legend, with lipid types represented with different colors. **(C)** Simulation system to study protein-protein cross-talks. Each system includes two prestin dimers at a 200-Å separation, embedded in a 400 Å × 200 Å bilayer, and with different relative orientations **(D)**. In each system, the two dimers are rotated around the membrane normal (z axis) in 45° increments. Given the two-fold symmetry of the protein, combinations of four different orientations per protein dimer (0°, 45°, 90°, and 135°, labeled as 1–4) cover all relative orientations of the two proteins at 45° intervals. Identical orientations are numbered the same, e.g., 45° and 225° are both labeled 2.

Three setups were constructed and simulated. In the first setup, we aimed to understand the impact of prestin on the overall deformation and lipid distribution of the membrane, for both contracted and expanded conformations. To enhance sampling of lipid-protein interactions and improve the statistics, this setup which we refer to as *quad-prestin* simulations, used four prestin dimers, separated by 200 Å, and placed in a large patch of lipid bilayer (400 × 400 Å^2^), using Insane^46^ (Fig. 1B).

In the second setup, we focused on the communication and cross-talk between the prestin dimers and how their individual deformation patterns might interfere with each other, and to compare their relative orientations. For this setup, referred to as *double-prestin* simulations, two prestin dimers, either in expanded or contracted conformations and separated by 200 Å, were placed in a rectangular lipid bilayer of dimensions 400 Å × 200 Å (Fig. 1C). To sample and compare multiple relative orientations, each prestin dimer was independently rotated around the *z* axis (membrane normal) at 45° intervals. Given the two-fold (C2) symmetry of prestin, placing each dimer at angles 0°, 45°, 90°, and 135° was sufficient to generate all possible 45° increments from 0 to 360° (Fig. 1D). To cover all the combinations of orientations of the two neighboring dimers, we constructed 16 different systems for each prestin’s conformational state (contracted or expanded) with the two protein dimers in one of the 4 different orientations: 4 orientations for Dimer I (0°, 45°, 90°, and 135°) × 4 orientations for Dimer II (0°, 45°, 90°, and 135°) = 16 systems. In these simulations, backbone restraints were used to maintain the positions and the relative orientations of the two dimers over the course of the simulations.

Finally, using a third, extended simulation, we monitored free rotation of two neighboring dimers with respect to each other. Using the *double-prestin* setup described above, with the two proteins both initially at 90°, a simulation was performed for 40 μs during which only the centers of masses of the dimers were restrained (to prevent their translation), but they were both allowed to freely rotate in their place. This simulation was used to sample the equilibrium distribution of the orientations.

All systems were then solvated and ionized with 150 mM NaCl using Insane,^46^ before the simulations.

### Simulation protocol

All systems were simulated with GROMACS,^47,48^ using the standard Martini v2.2 simulation settings.^49^ A 20-fs timestep was employed, and the temperature was maintained at 310 K with the velocity-rescaling thermostat^50^ using a coupling time constant of 1 ps. A semi-isotropic, 1-bar pressure was maintained using the Berendsen barostat^51^ with a compressibility of 3×10^-4^ bar and a relaxation time constant of 5 ps. Initially, the systems were energy minimized for 1,000 steps, followed by short equilibration runs (18 ns) while lipid headgroups and protein backbones were restrained. Over this initial period, the restraints on lipid headgroups were gradually decreased in several steps from *k* = 200kJ.mol^-1^.nm^-2^ to zero), whereas the protein backbone restraints (*k* = 1000kJ.mol^-1^.nm^-2^) were maintained. For the production runs, the *quad-prestin* systems were simulated for 20 μs each (Table S1). The 16 *double-prestin* systems with different orientations were simulated for 10μs each (Table S2). The third final simulation system was initiated from the *double-prestin* system with both dimers at 90° (parallel) after 10 μs of equilibration. After removing the backbone restraints to allow the system to rotate freely, and introducing center of mass restraints for each prestin dimer to prevent their translation, the system was simulated for 40 μs.

### Analysis

VMD^44^ was used as a main analysis environment and to generate all the molecular images. Inhouse scripts were developed for specific geometrical analyses of the simulated systems, as described below. The position of a lipid or its distance to the protein was evaluated by the CG bead representing its phosphodiester PO_4_ in the case of phospholipids and sphingomyelin, or the hydroxy group in the case of cholesterol.

#### Membrane deformation

The membrane deformation induced by prestin was evaluated by the height of the lipids (separation from the bilayer midplane, referred to as lipid elevation or depression) in each leaflet. The *z* position (along the membrane normal) of the phosphate beads (PO_4_ beads in Martini) of phospholipids was used for this purpose, with the bilayer midplane at *z* = 0. The data, averaged over the last 5 *μ*s of each trajectory, were then binned in 2 × 2 Å^2^ bins to generate a 2-dimensional histogram in the *xy* plane (membrane plane). These histograms represent the average height (thickness) of each leaflet at different (*x, y*) coordinates. To quantify the propagation of membrane deformation, the maximum and minuimum lipid heights were recorded at each distance (radius) with regard to the protein’s center in each frame, and their time-averaged values were plotted as a function of radius.

#### Lipid distribution around prestin

To evaluate the effect of prestin on lipid distribution in the membrane, lipid counts within 7 Å of the protein were averaged for each lipid type over the 20 μs of each *quad-prestin* simulation and compared to their bulk density. A depletion-enrichment index for lipid type L was then calculated as the ratio of the local and global fractions of the lipid:^42^

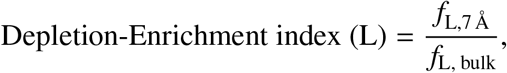

where

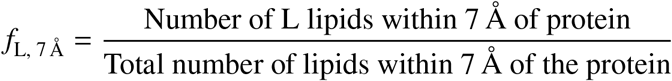

and

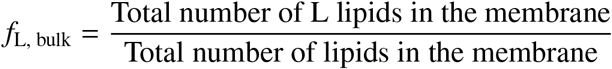

#### Membrane bending modulus

To calculate the bending modulus, we used the Helfrich-Canham (HC) theory, which relates the equilibrium fluctuations of a membrane to its elastic properties.^52–54^ The fluctuations in the *z* direction were calculated for the phosphate bead (PO_4_) of each phospholipid and then histogrammed over the membrane (*xy*) plane using 1 × 1 Å^2^ bins. The fluctuations were then transformed to the Fourier space and employed in the following equation to calculate the bending modulus:^55^

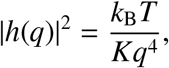

where *h*(*q*) is fluctuation spectrum in the *z* direction in the Fourier space, *k_B_* is Boltzmann constant, *T* is the temperature, *K* is the bending modulus, and *q* is the magnitude of the wavevector (i.e., the −1 wavenumber, Å^-1^).^55^

To obtain the bending modulus, *K*, |*h*(*q*)|^2^*q*^4^ was plotted with respect to *q*, and then fitted to a line parallel to the *x* axis. The fitting was performed for low *q* values, which approximately form a line and represent the most dominant mode shapes in the membrane bending.^55^ The *y* value of the fitted line is equivalent to 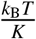 and can be used to calculate *K*. All the *K* values were divided by the maximum *K* for each prestin conformation, generating scaled bending moduli, which were used for comparing the systems.

## Results and Discussion

### Prestin induces drastic, anisotropic deformation in the membrane

To investigate the effect of prestin on the surrounding membrane, we probed the structure of a membrane composed of 4 prestin dimers (separated by 200 Å) embedded in a lipid mixture (quad-prestin) over 20 μs of CG MD simulation (Fig. 1B). The membrane structure was analyzed using the *z* positions of lipid headgroups in small bins to provide a metric with sufficiently detailed spatial resolution. The constructed 2D histograms indicates drastic, yet non-uniform deformation of the membrane by prestin, manifested by elevation or depression of the lipids flanking different sides of the protein, in both the inner and outer leaflets (Fig. 2A-B). The membrane was centered at *z* = 0, with the phospholipid headgroups of the outer and inner leaflets initially positioned at approximately *z* = +20 and *z* = −20 Å, respectively. During the simulation, the phospholipid headgroups displace to *z* values ranging from +5 to +30 Å in the outer leaflet, and from −5 to −30 Å in the inner leaflet, corresponding to displacements as large as 15 A from their initial positions in each leaflet (Fig. 2A-B).

**Figure 2:**
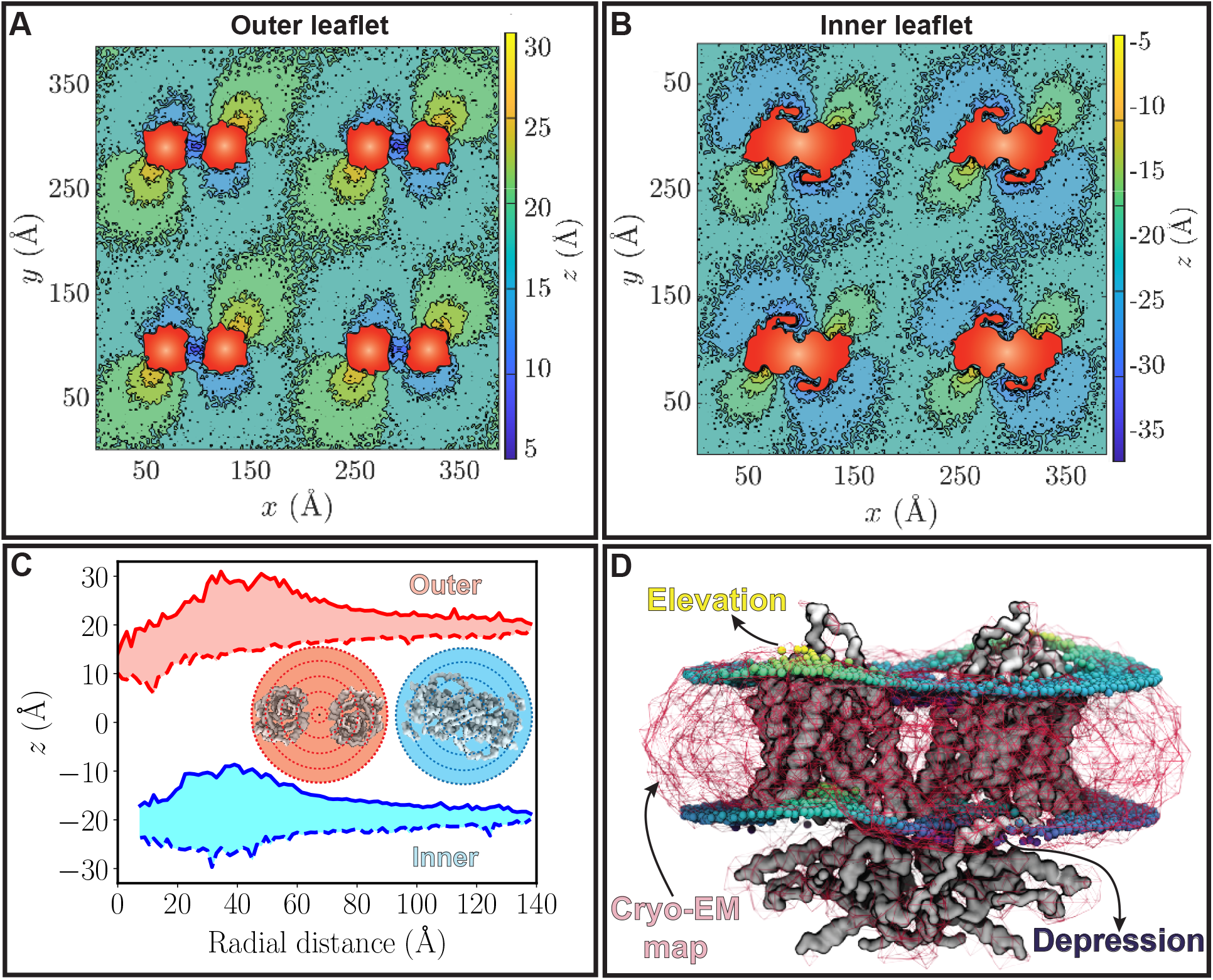
Prestin-induced, non-uniform membrane deformation. **(A)** and **(B)** 2D histograms of the *z* positions of lipid headgroups for outer and inner leaflets. Both leaflets are viewed from the extracellular side. Histograms are constructed from the last 5μs of 20-μs trajectories. The cross sectional areas of the prestin dimers in each leaflet are colored in red. **(C)** range of *z* values of lipid headgroups in radial sections (drawn in the inset) around the protein, red for the outer and blue for the inner leaflet. The maximum elevation values are represented with solid and minimum values with dashed lines, whereas the range of observed *z* values is shown in light red and blue. The blue curves do not start from zero, since in the inner leaflet there are no lipids in that region. **(D)** Agreement between the lipid density in the cryo-EM map and the phospholipid distribution from the simulations, averaged over the last 5 *μ*s of one of the trajectories. The cryo-EM density is represented as a red mesh. The coloring of headgroups is based on their *z* values, with highest and lowest *z* shown in yellow and blue, respectively. Elevated and depressed regions of the membrane form MD simulations overlap with corresponding areas in the cryo-EM map.

The depressed lipids in the outer leaflet are primarily associated with the core domains, particularly in the region between the two prestin protomers, highlighted by *z* values as low as *z* = 5 Å (Fig. 2A and S2A). Notably, the region between the two protomers was initially free of lipids but becomes completely filled during the simulation in all four protein copies (Fig. S2A). In contrast to the depressed membrane regions, elevated lipids in the outer leaflet arise in the vicinity of the gate domains of prestin where lipid elevations as large as *z* = 30 Å are observed.

Similar to the outer leaflet, the pattern of membrane deformation in the inner leaflet can be characterized by lipid elevation in the proximity of the gate domains and lipid depression in the vicinity of the core domains (Fig. 2B and S2B). In the inner leaflet, the elevated lipids reach *z* values of about −5 Å, whereas in the depressed region lipids sink to as low as *z* = −30 Å (Fig. 2B). Similar membrane deformation patterns were observed for both the outer and inner leaflets in the case of the expanded prestin conformation (Fig. S3A-B).

In order to quantify the range over which prestin deforms the membrane, we defined radial regions around the protein, and for each region, the minimum and maximum *z* values of lipids were recorded. The analysis clearly establishes the long range of prestin-induced membrane deformation, extending as long as 10 nm away from the protein(Figs. 2A-C and S3A-B). Membrane deformation around the protein follows the two-fold symmetry of prestin’s dimeric structures (Figs. 2A-B and S3A-B), and deformation patterns of individual prestin dimers are highly consistent, supporting the convergence of the simulations (Figs. 2A-B and S3A-B).

The deformation patterns captured here indicate a strong, but heterogeneous membrane response to prestin and are in close agreement with the observed lipid densities around the protein in cryoEM studies of nanodisc-embedded prestin.^18,19^ Overlapping the cryo-EM map for prestin structure (e.g., in the contracted conformation (PDB ID: 7LGW)^18^) and the positions of phospholipid headgroups from the simulations, a close agreement between the computational and experimental results is evident (Fig. 2D).

### Prestin alignment mediated by membrane deformation

To characterize membrane deformation patterns produced by different prestin arrangements, we designed an array of of simulation systems each with two prestin dimers embedded in a rectangular membrane at a fixed prestin-prestin distance but with different relative orientations (Fig. 3). Membrane deformation patterns were monitored by constructing lipid height heatmaps (Fig. 3A-B for contracted prestin, and Fig. S4A-B for expanded prestin). The heatmpas immediately highlight an important relationship of membrane deformation patterns, namely the significant overlap between the patterns induced by neighboring prestin dimers, indicating that the proteins might indirectly influence each other’s orientation through their effects on the membrane. For example, the configuration with two prestin dimers at 0° ([0°, 0°]; top left in Fig. 3A-B) generates non-matching, destructive deformation patterns (Fig. 3A-B). However, as the prestin dimers are rotated in other simulation systems, the membrane deformation patterns start to align with each other, thereby creating constructive interference profiles, e.g., in the [45°, 45°] configuration (Fig. 3A-B). Interestingly, when the membrane deformation patterns are constructively aligned, e.g., in the [45°, 45°] configuration, the structural effect of prestin on the membrane seems to be propagating over a longer range (Fig. 3A-B). The membrane deformation patterns for expanded prestin dimers are similar to those obtained from contracted systems, but generally with a smaller propagation range (Figs. 3A-B and S4A-B).

**Figure 3:**
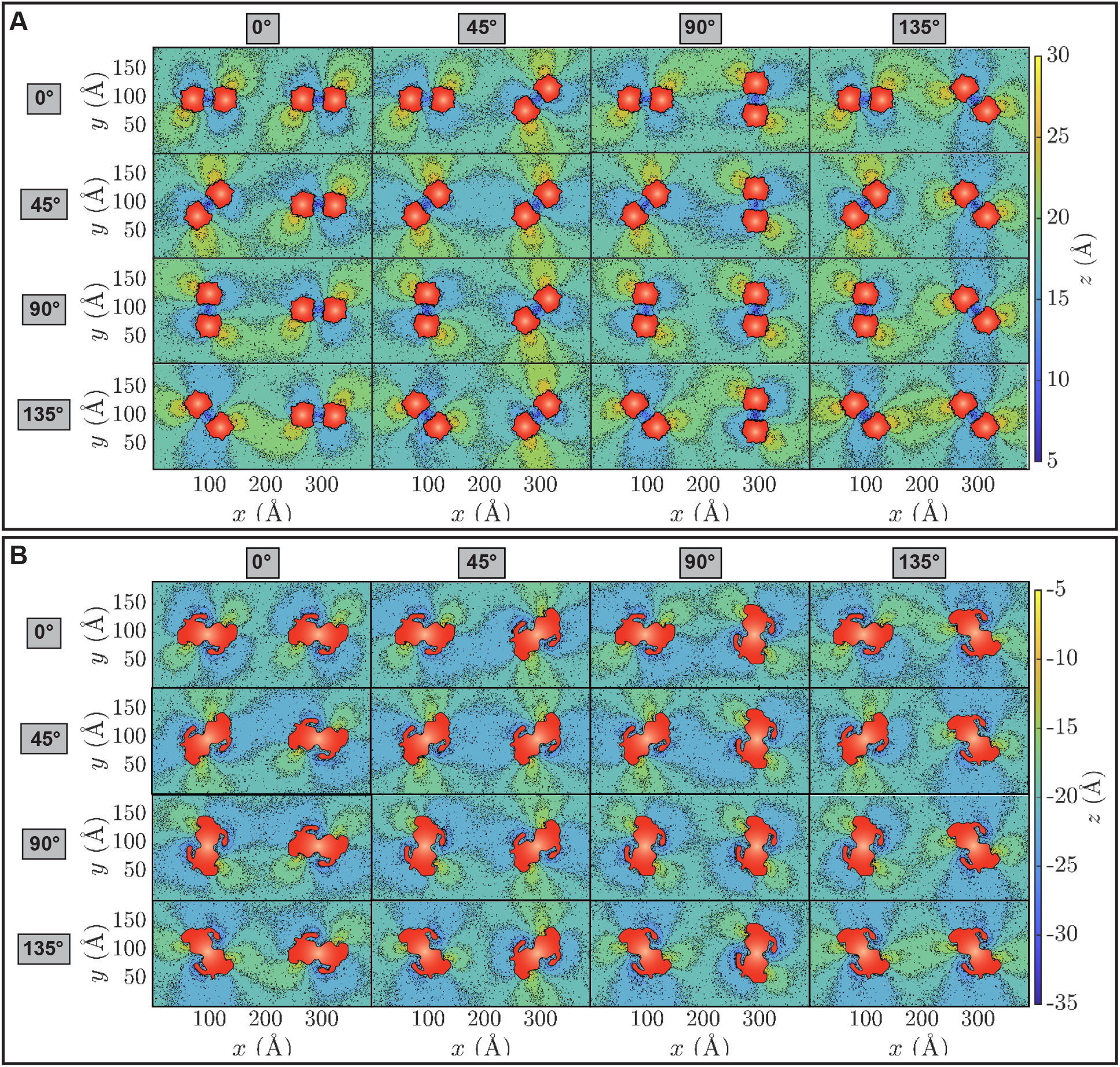
Constructive or destructive interference of deformation patterns induced by prestin’s different orientations. Heatmaps of phospholipid height in the outer **(A)** and inner **(B)** leaflets are calculated using the procedure described in Fig. 2. The prestin dimers are placed at (*x, y*) = (100,100) Å and (300,100) Å, respectively. The orientation of Dimer I (angle with respect to the *x* axis) is specified on the left, and for Dimer II on the top of each panel. Given the symmetry relation between prestin’s protomers, four angles (0°, 45°, 90°, and 135°) for each prestin dimer are sufficient to cover all possible orientations (at 45° intervals), resulting in 16 different combinations of two dimers. The protein cross sectional area in each leaflet is drawn in red.

To quantitatively compare the different deformation patterns caused by prestin’s orientation and their effect on the membrane structural response, the associated membrane stiffness was estimated by calculating the membrane bending modulus, which is directly proportional to membrane bending energy. The results show that when membrane deformation patterns are in line, the membrane bending modulus is minimal (Fig. 4A-B), suggesting that in these configurations the change in the cellular membrane shape in response to the mechanical activity of the protein might be maximal. In contrast, in cases with destructive interference between the deformation patterns, i.e., when depressed lipids around one prestin dimer face lipids elevated by a neighboring dimer, the membrane exhibits a larger degree of rigidity (Figs. 3A-B and 4A-B).

**Figure 4:**
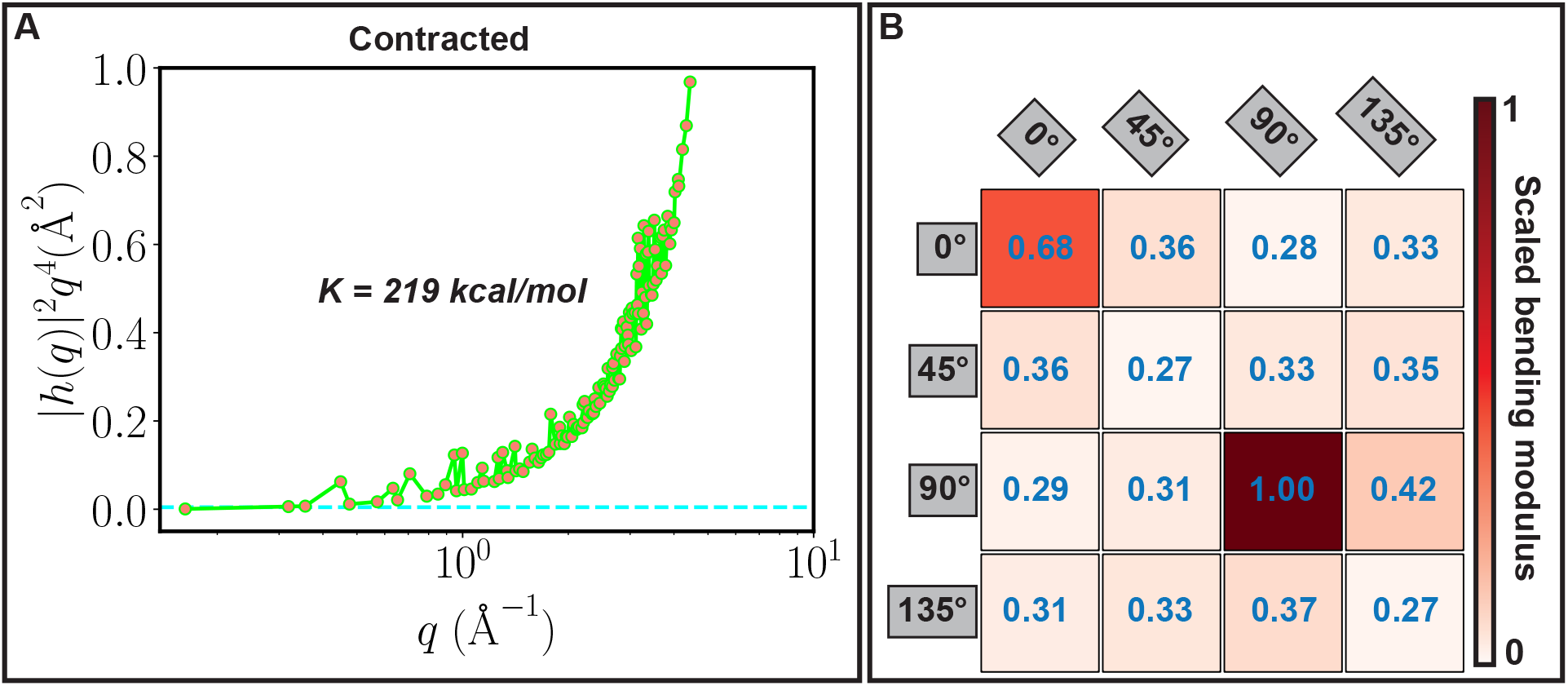
Membrane bending moduli calculated for different prestin arrangements. **(A)** |*h*(*q*)|^2^*q*^4^ plotted vs. *q*, wavenumber, for the system with two prestin dimers both at 0° ([0°, 0°]). The y value of the horizontal line fitted to the function at low *q* values (blue dashed line) is equivalent to 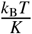, and is used to calculate *K*. For the represented system here, *K* = 219kcal/mol. **(B)** Bending moduli for the membranes with two prestin dimers in different relative orientations (Fig. 3), normalized by the maximum value for the [90°, 90°] system. The orientation of Dimer 1, placed at (*x, y*) = (100, 100) Å is shown on the left, and for Dimer II, placed at (*x, y*) = (300, 100) Å, on the top of the table. The color shade in each box also represents the strength of the scaled bending modulus, and defined in the scale bar. The minimum bending moduli are captured when the two prestin dimers are both at either 45° or 135°.

The lowest bending modulus (softest membrane) for contracted prestin simulations is obtained when the dimers are both at either 45° or 135°. In these configurations, the depressed and elevated lipids surrounding reach their maximum propagation to fully extend between the two prestin dimers (Figs. 3A-B and S4A-B). Note that systems with both prestin dimers either at 45° or at 135° correspond to the same relative configurations (due to the C2 symmetry in prestin’s dimer). For the same system, the highest bending modulus is obtained when the neighboring dimers are both at 90° ([90°, 90°]; Fig. 4A-B), where maximal mismatch between depressed and elevated lipids from the two dimers arises. Similar profiles are obtained for the simulations of expanded prestin (Fig. S5A-B). The lowest bending moduli for both contracted and expanded prestin correspond to [45°, 45°] or [135°, 135°] orientations, although for the contracted system most bending moduli remain close to their lowest value (bright shades; Fig. 3B), whereas in expanded prestin they differ significantly (dark shades; Fig. S4B). One might speculate that in its contracted form prestin is more free to alter its relative orientation, whereas in its expanded form it is more confined to a particular orientation.

The calculated membrane deformation patterns suggest that prestin dimers on the surface of OHCs might be organized with a preferred orientation of 45/135°, such that their elevated and depressed lipid regions align, thereby achieving the longest range and largest degree of membrane deformation (Figs. 3A-B and S4A-B). Given the prestin density in the simulations, its favored relative orientations observed here are of direct relevance to the crowded arrangement of prestin in the OHCs.

To further validate the preferred prestin-prestin configuration in the membrane, we designed another simulation in which neighboring prestin dimers were allowed to freely rotate (Fig. 5A). The orientations of the two dimers (θ_1_ and θ_2_) were then monitored during a 40-μs simulation (Fig. 5B). The time evolution of the angles confirms our conclusion regarding the preferred orientations of prestin in the membrane: either at ~45° (same as 225°) or at ~135° (same as 315°) (Fig. 5B and Supplementary Movie). Furthermore, the neighboring dimers seem to sense and follow each other’s orientation, in that when one of them changes its orientation, the other follows. For instance, at *t* =~20μs, when one dimer alters its orientation from ~225° to ~135° (orange trace in Fig. 5B and Supplementary Movie), the second dimer (green trace) follows immediately and changes its orientation from ~225° to ~315° (Fig. 5B). Such membrane-mediated communication between prestin dimers is evident at several time points during the simulation (Fig. 5B and Supplementary Movie). The free energy landscape in θ_1_/θ_2_ phase space, constructed from the last 30 μs of the simulation, reveals three different minima (Fig. 5C), all close to the optimal configurations predicted from our earlier simulations, i.e., corresponding to [45°, 45°] and [135°, 135°] configurations (Fig. 5C).

**Figure 5:**
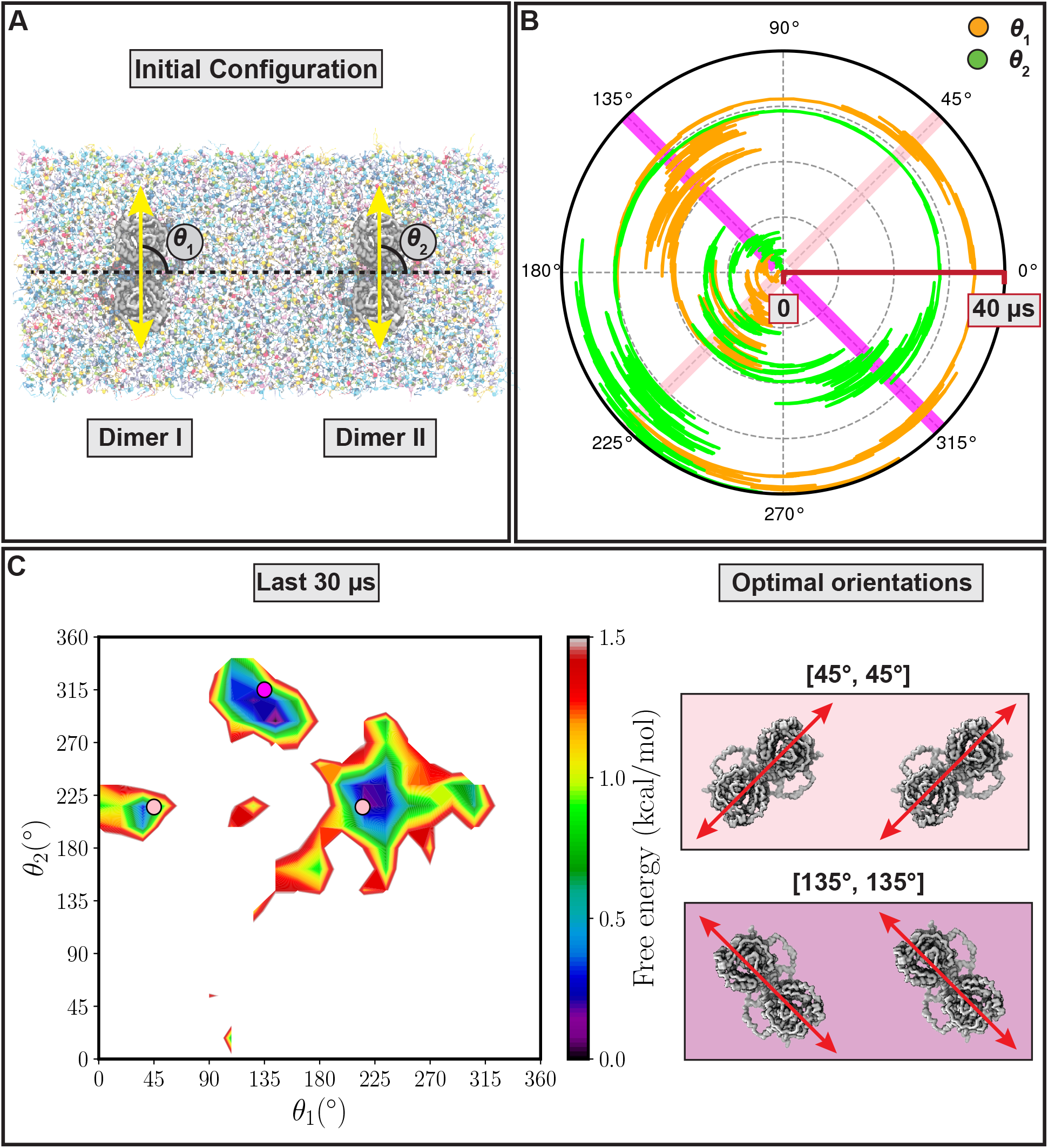
Prestin-prestin communication through the membrane. **(A)** Initial system setup with two contracted prestin dimers at 90° [90°, 90°]. The center of mass of each dimer is restrained, but they are allowed to rotate freely. Orientation of the two prestin dimers are denoted as *θ*_1_ and *θ*_2_, respectively. **(B)** Time series of *θ*_1_ (orange) and *θ*_2_ (green) during the 40 μs simulation. The radial direction in the polar plot represents the time from 0 to 40 μs. See also Supplementary Movie. **(C)** Left: Free energy landscape in *θ*_1_/*θ*_2_ space, constructed from the last 30 μs of the trajectory. The landscape consists of three main minima. The pink and magenta circles correspond to [45°, 45°] and [135°, 135°] prestin-prestin configurations, respectively, as shown in the Right side of the panel, and also highlighted in **B** by diagonal bars.

### Enrichment of PI and CHOL around prestin

In addition to membrane deformation, we also observe differential enrichment or depletion of different lipid types around prestin (Fig. 6A). For CHOL, and particularly for PI, the simulations record large increases in their local density around the protein (Depletion-Enrichment indices of 1.4 and 2.2, respectively; Fig. 6B). As shown in Fig. 6A, the initial numbers of 4 PI and 8 CHOL lipids in the vicinity of prestin in each leaflet increase to 8-15 PI and 10-15 CHOL per leaflet over the 20-μs course of the simulation. In contrast, the number of PC, PE, PG, and SM lipids in the vicinity of the protein in both the outer and inner leaflets slightly drops, mostly during the first 2 μs (Depletion-Enrichment indices less than one; Fig. 6B). Despite being a charged lipid, the PS count close to prestin remains relatively unaltered during the simulation (Depletion-Enrichment index of nearly one; Fig. 6B). Approximately similar counts and enrichment/depletion indices are obtained for expanded prestin simulations (Fig. S6A-B).

**Figure 6:**
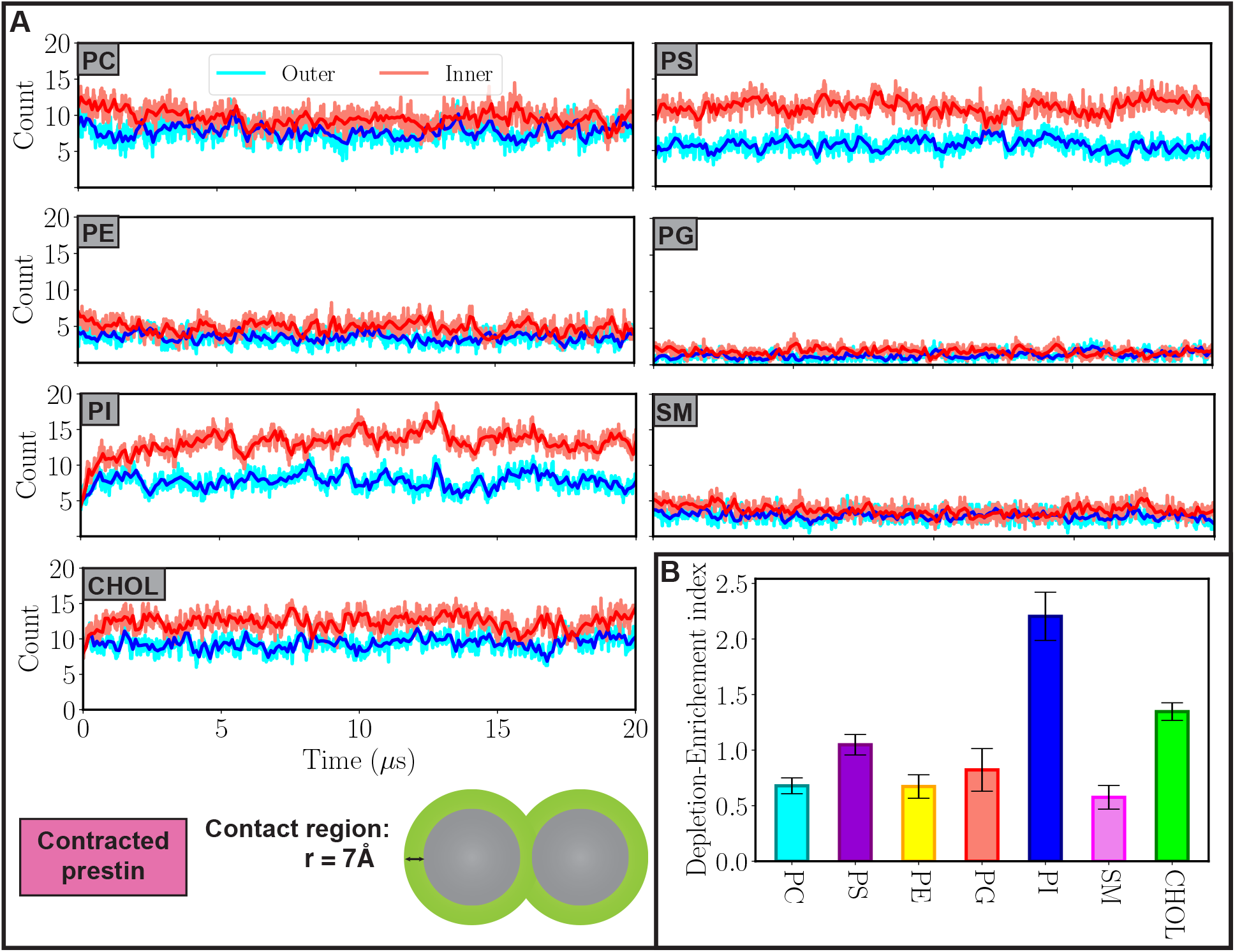
Enrichment/depletion of different lipids around contracted prestin. **(A)** Time series of the number of different lipid types within 7 Å of a prestin dimer, averaged over four dimers, in the outer (blue) and inner (red) leaflets. PI lipids accumulate the most around prestin in the inner leaflet. **(B)** Depletion-Enrichment index of each lipid type (the ratio of the local and bulk fractions of the lipid). PI and CHOL show the highest enrichment, whereas SM and PE show the largest degree of depletion.

To pinpoint high-density regions of PI and CHOL around prestin, we calculated the 2D distribution of these lipids during the last 5 μs of the trajectory for all four prestin dimers in the quad-prestin simulation. PI accumulation is mostly observed around the core and gate domains, with higher densities around the latter in the outer leaflet, and around the former in the inner leaflet (Fig. 7A). In the case of CHOL, high-occupancy regions are observed more scattered around the protein (Fig. 7A). Of particular significance, two CHOL high-density regions are positioned symmetrically in between the two protomers in the outer leaflet (Fig. 7A), i.e., at the interface of the core and gate domains of each protomer (Fig. 7A). The MD trajectory shows that, once bound, these inter-dimeric CHOL molecules tend to stay in this site while sampling different conformations (see snapshots from *t* = 10 – 20μs in Fig. 7B). Supporting these observations, the cryo-EM model of contracted prestin contains two symmetrically positioned CHOL molecules at this location between the two protomers.^18^ Despite thermally driven fluctuations, the CHOL trajectory samples conformations similar to cryo-EM poses (Fig. 7B). Interestingly, the region between the two prestin protomers remains free of PI throughout the simulation (Fig. 7A).

**Figure 7:**
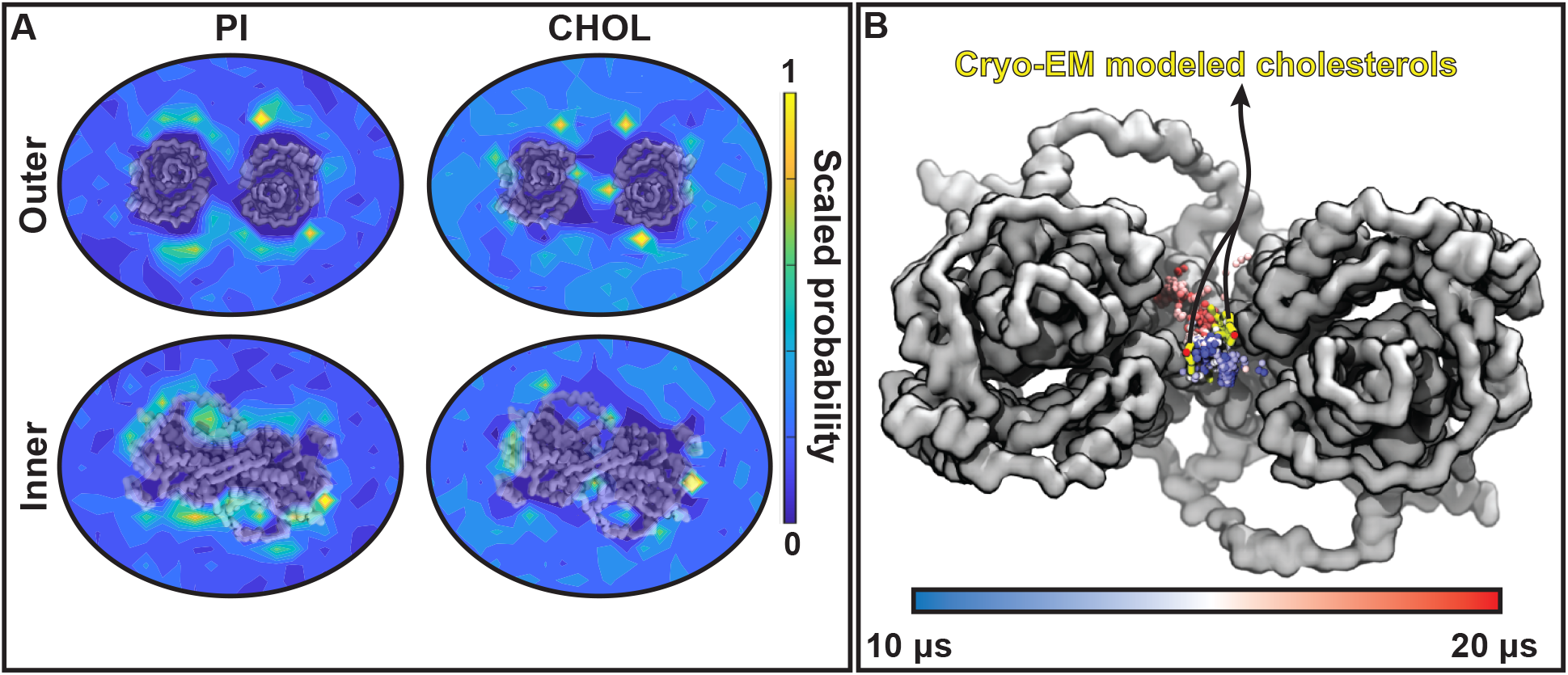
Accumulation of PI and CHOL around prestin. **(A)** Heatmaps representing the distribution of PI and CHOL over the last 5*μs* of the quad-prestin trajectory in the contracted conformation. Both PI and CHOL cluster closely to the core and gate domains of prestin. The region in between the two protomers contains two distinct CHOL binding sites but remains free of PI during the simulation. **(B)** Comparison of cryo-EM and MD CHOL. The trajectory of one of the CHOL molecules in between the two protomers during the last 10 μs of the simulation is shown in varying colors representing the time (scale bar). Only the hydroxy bead is shown for each frame. The cryo-EM CHOL models are shown in yellow.

Clustering of PI and CHOL has also been reported for other anionic carriers. In a MD study of Band 3 (*aka* AE1 or SLC4A1), a Cl^-^/HCO^-^ exchanger in red blood cells and kidneys, annular lipids were found to be enriched in PI and CHOL. ^56^ Clustering of PI and CHOL lipids close to prestin, reported in our study, might play a role in prestin’s conformational dynamics, as these lipids can bind membrane proteins through specific, often functionally important, binding sites.^57–60^

### Concluding Remarks

Prestin is a high-density motor protein on the surface of the OHCs, whose conformational changes constitute the main mechanism of cochlear sound amplification in mammalian auditory systems. Recent advances in structural studies have characterized the protein in both active and inactive conformational states. However, the extent and nature of its interaction with the lipids and the ensuing membrane deformation by prestin, which are at the heart of its effect on the shape of the OHCs remained elusive.

Here, using an extensive set of Martini CG MD simulations (collectively, 520 μs) we characterize the nature of membrane structural effects of prestin. The simulations reveal drastic, anisotropic membrane deformation patterns, manifested by regions with elevated or depressed lipids, induced by the gate and core domains of protein. These deformation patterns are long-range and last for ~10nm in both the outer and inner leaflets. The elevated and depressed regions are enriched in CHOL and PI, suggesting possible effects on the function of prestin. The observed drastic membrane deformation and its long range of propagation suggested possible, lipid-mediated communication between prestin dimers at surface densities observed in OHCs. This aspect was investigated by simulations involving pairs of prestin dimers with different relative orientations and quantifying their effects on membrane structure and rigidity. Both rotationally restrained and free-rotating simulations clearly highlight favorable orientations of neighboring prestin molecules that allow for maximal alignment of deformation patterns (Fig. 8) and a reduction in the bending modulus of the membrane, thereby resulting in a more responsive membrane to mechanical forces caused by prestin. These structural and mechanical properties of the membrane are of high relevance to prestin’s biological role as the main element of cochlear sound amplification.

**Figure 8:**
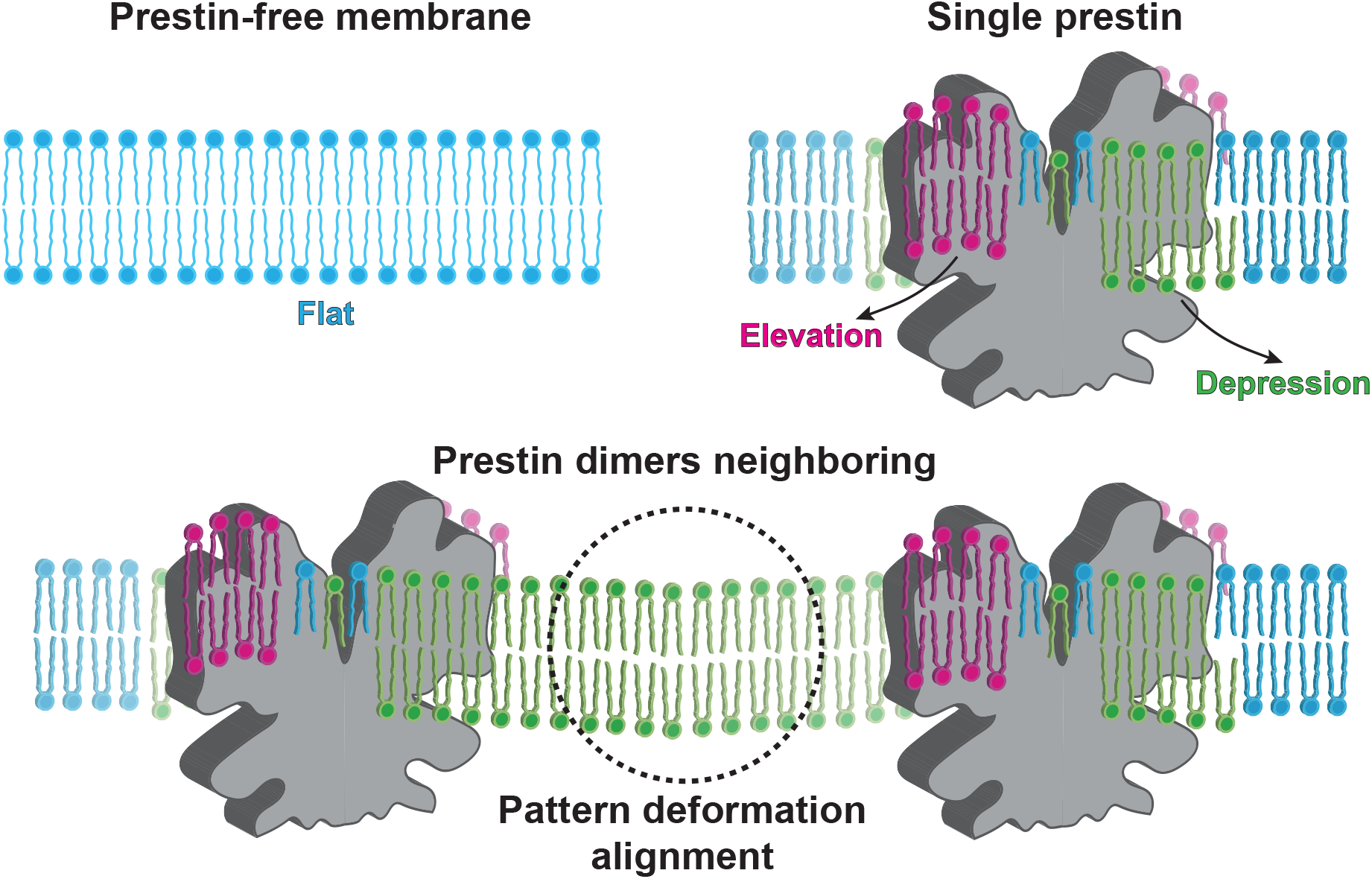
Schematic representation of membrane deformation patterns induced by a single or multiple prestin dimers. A prestin-free membrane remains flat, as highlighted by the even height of its lipids (blue). Prestin deforms the membrane anisotropically by elevating or depressing lipids (magenta and green, respectively) in its vicinity. In a multi-protein arrangement, as in the OHCs, membrane deformation patterns of individual prestin dimers can align constructively, with matching elevated and depressed lipids. These favorable prestin-prestin configurations, which likely represent its preferred arrangement in the OHCs, offer a longer range of membrane deformation as well as a softer membrane with a greater extent of response to prestin’s mechanical role in sound amplification.

## Supporting information

Animation displaying alignment of prestin neighbors in the membrane

## Acknowledgement

The authors acknowlede stimulating conversations and discussion with Drs. E. Gouaux, J. Ge, and J. Elferich at Vollum Institute, Oregon Health & Science University. This research is supported by National Institutes of Health grants R01-GM123455 and P41-GM104601 (to E.T.). Simulations in this study have been performed using allocations at National Science Foundation Supercomputing Centers (XSEDE grant number MCA06N060), and the Blue Waters Petascale Computing Facility of National Center for Supercomputing Applications (NCSA) at University of Illinois at Urbana-Champaign, which is supported by the National Science Foundation (awards OCI-0725070 and ACI-1238993) and the State of Illinois.

## References

[1] Martin L Basch, Rogers M Brown, Hsin-I Jen, and Andrew K Groves. Where hearing starts: the development of the mammalian cochlea. Journal of Anatomy, 228(2):233–254, 2016.

[2] Bruce Masterton, Henry Heffner, and Richard Ravizza. The evolution of human hearing. The Journal of the Acoustical Society of America, 45(4):966–985, 1969.

[3] Jonathan Ashmore. Outer hair cells and electromotility. Cold Spring Harbor Perspectives in Medicine, 9(7):a033522, 2019.

[4] William E Brownell, Charles R Bader, Daniel Bertrand, and Yves De Ribaupierre. Evoked mechanical responses of isolated cochlear outer hair cells. Science, 227(4683):194–196, 1985.

[5] JF Ashmore. A fast motile response in guinea-pig outer hair cells: the cellular basis of the cochlear amplifier. The Journal of Physiology, 388(1):323–347, 1987.

[6] JE Gale and JF Ashmore. The outer hair cell motor in membrane patches. Pflügers Archiv – European Journal of Physiology, 434(3):267–271, 1997.

[7] Stuart L Johnson, Maryline Beurg, Walter Marcotti, and Robert Fettiplace. Prestin-driven cochlear amplification is not limited by the outer hair cell membrane time constant. Neuron, 70(6):1143–1154, 2011.

[8] David ZZ He, Sándor Lovas, Yu Ai, Yi Li, and Kirk W Beisel. Prestin at year 14: progress and prospect. Hearing Research, 311:25–35, 2014.

[9] Peter Dallos, Xudong Wu, Mary Ann Cheatham, Jiangang Gao, Jing Zheng, Charles T Anderson, Shuping Jia, Xiang Wang, Wendy HY Cheng, Soma Sengupta, et al. Prestin-based outer hair cell motility is necessary for mammalian cochlear amplification. Neuron, 58(3):333–339, 2008.

[10] Anthony W Gummer and Serena Preyer. Cochlear amplification and its pathology: Emphasis on the role of the tectorial membrane. Ear, Nose & Throat Journal, 76(3):151–163, 1997.

[11] Joseph Santos-Sacchi, Dhasakumar Navaratnam, Rob Raphael, and Dominik Oliver. Prestin: molecular mechanisms underlying outer hair cell electromotility. In Understanding the Cochlea, pages 113–145. Springer, 2017.

[12] Xudong Wu, Jiangang Gao, Yunkai Guo, and Jian Zuo. Hearing threshold elevation precedes hair-cell loss in prestin knockout mice. Molecular Brain Research, 126(1):30–37, 2004.

[13] Hannes Lohi, Minna Kujala, Erja Kerkelä, Ulpu Saarialho-Kere, Marjo Kestilä, and Juha Kere. Mapping of five new putative anion transporter genes in human and characterization of slc26a6, a candidate gene for pancreatic anion exchanger. Genomics, 70(1):102–112, 2000.

[14] Silvia Detro-Dassen, Michael Schanzler, Heike Lauks, Ina Martin, Sonja Meyer zu Berstenhorst, Doreen Nothmann, Delany Torres-Salazar, Patricia Hidalgo, Gunther Schmalzing, and Christoph Fahlke. Conserved dimeric subunit stoichiometry of SLC26 multifunctional anion exchangers. Journal of Biological Chemistry, 283(7):4177–4188, 2008.

[15] Dominik Oliver, David ZZ He, Nikolaj Klöcker, Jost Ludwig, Uwe Schulte, Siegfried Waldegger, JP Ruppersberg, Peter Dallos, and Bernd Fakler. Intracellular anions as the voltage sensor of prestin, the outer hair cell motor protein. Science, 292(5525):2340–2343, 2001.

[16] David ZZ He, Burt N Evans, and Peter Dallos. First appearance and development of electro-motility in neonatal gerbil outer hair cells. Hearing Research, 78(1):77–90, 1994.

[17] Jing Zheng, Weixing Shen, David ZZ He, Kevin B Long, Laird D Madison, and Peter Dallos. Prestin is the motor protein of cochlear outer hair cells. Nature, 405(6783):149–155, 2000.

[18] Jingpeng Ge, Johannes Elferich, Sepehr Dehghani-Ghahnaviyeh, Zhiyu Zhao, Marc Meadows, Henrique von Gersdorff, Emad Tajkhorshid, and Eric Gouaux. Molecular mechanism of prestin electromotive signal amplification. Cell, 184(18):4669–4679, 2021.

[19] Navid Bavi, Michael David Clark, Gustavo F Contreras, Rong Shen, Bharat G Reddy, Wieslawa Milewski, and Eduardo Perozo. Prestin’s conformational cycle underlies outer hair cell electromotility. Nature, pages 1–11, 2021.

[20] Dmitry Gorbunov, Mattia Sturlese, Florian Nies, Murielle Kluge, Massimo Bellanda, Roberto Battistutta, and Dominik Oliver. Molecular architecture and the structural basis for anion interaction in prestin and SLC26 transporters. Nature Communications, 5(1):1–13, 2014.

[21] Yung-Ning Chang, Eva A Jaumann, Katrin Reichel, Julia Hartmann, Dominik Oliver, Gerhard Hummer, Benesh Joseph, and Eric R Geertsma. Structural basis for functional interactions in dimers of SLC26 transporters. Nature Communications, 10(1):1–10, 2019.

[22] Federico Kalinec, Matthew C Holley, Kuni H Iwasa, David J Lim, and Bechara Kachar. A membrane-based force generation mechanism in auditory sensory cells. Proceedings of the National Academy of Sciences, USA, 89(18):8671–8675, 1992.

[23] Peter Dallos and Bernd Fakler. Prestin, a new type of motor protein. Nature Reviews Molecular Cell Biology, 3(2):104–111, 2002.

[24] KH Iwasa. Effect of stress on the membrane capacitance of the auditory outer hair cell. Biophysical Journal, 65(1):492–498, 1993.

[25] Joseph Santos-Sacchi, Weixing Shen, Jing Zheng, and Peter Dallos. Effects of membrane potential and tension on prestin, the outer hair cell lateral membrane motor protein. The Journal of Physiology, 531(3):661–666, 2001.

[26] David ZZ He, Shuping Jia, and Peter Dallos. Prestin and the dynamic stiffness of cochlear outer hair cells. Journal of Neuroscience, 23(27):9089–9096, 2003.

[27] Enrique G Navarrete and Joseph Santos-Sacchi. On the effect of prestin on the electrical breakdown of cell membranes. Biophysical Journal, 90(3):967–974, 2006.

[28] Rui Zhang, Feng Qian, Lavanya Rajagopalan, Fred A Pereira, William E Brownell, and Bahman Anvari. Prestin modulates mechanics and electromechanical force of the plasma membrane. Biophysical Journal, 93(1):L07–L09, 2007.

[29] Lavanya Rajagopalan, Jennifer N Greeson, Anping Xia, Haiying Liu, Angela Sturm, Robert M Raphael, Amy L Davidson, John S Oghalai, Fred A Pereira, and William E Brownell. Tuning of the outer hair cell motor by membrane cholesterol. Journal of Biological Chemistry, 282(50):36659–36670, 2007.

[30] RI Kamar, LE Organ-Darling, and RM Raphael. Membrane cholesterol strongly influences confined diffusion of prestin. Biophysical Journal, 103(8):1627–1636, 2012.

[31] AG Lee. Lipid–protein interactions in biological membranes: a structural perspective. Biochimica et Biophysica Acta – Biomembranes, 1612(1):1–40, 2003.

[32] Nelson P Barrera, Min Zhou, and Carol V Robinson. The role of lipids in defining membrane protein interactions: insights from mass spectrometry. Trends in Cell Biology, 23(1):1–8, 2013.

[33] Melanie P. Muller, Tao Jiang, Chang Sun, Muyun Lihan, Shashank Pant, Paween Mahinthichaichan, Anda Trifan, and Emad Tajkhorshid. Characterization of lipid-protein interactions and lipid-mediated modulation of membrane protein function through molecular simulations. Chemical Reviews, 119:6086–6161, 2019.

[34] Nicolas Sapay and D Peter Tieleman. Molecular dynamics simulation of lipid–protein interactions. Current Topics in Membranes, 60:111–130, 2008.

[35] Sundeep S. Deol, Peter J. Bond, Carmen Domene, and Mark S.P. Sansom. Lipid-protein interactions of integral membrane proteins: A comparative study. Biophysical Journal, 87:3737–3749, 2004.

[36] Michael C. Pitman, Alan Grossfield, Frank Suits, and Scott E. Feller. Role of cholesterol and polyunsaturated chains in lipid-protein interactions: Molecular dynamics simulation of rhodopsin in a realistic membrane environment. Journal of the American Chemical Society, 127(13):4576–4577, 2005.

[37] Sepehr Dehghani-Ghahnaviyeh, Karan Kapoor, and Emad Tajkhorshid. Conformational changes in the nucleotide-binding domains of P-glycoprotein induced by ATP hydrolysis. FEBS Letters, 595(6):735–749, 2021.

[38] Clara Abaurrea Velasco, Sepehr Dehghani Ghahnaviyeh, Hossein Nejat Pishkenari, Thorsten Auth, and Gerhard Gompper. Complex self-propelled rings: a minimal model for cell motility. Soft Matter, 13(35):5865–5876, 2017.

[39] E. Lindahl and M. S. P. Sansom. Membrane proteins: molecular dynamics simulations. Current Opinion in Structural Biology, 18:425–431, 2008.

[40] DP Tieleman, BI Sejdiu, EA Cino, P Smith, E Barreto-Ojeda, HM Khan, and V Corradi. Insights into lipid-protein interactions from computer simulations. Biophysical Reviews, pages 1–9, 2021.

[41] Valentina Corradi, Besian I. Sejdiu, Haydee Mesa-Galloso, Haleh Abdizadeh, Sergei Yu Noskov, Siewert J. Marrink, and D. Peter Tieleman. Emerging diversity in lipid-protein interactions. Chemical Reviews, 119(9):5775–5848, 2019.

[42] Valentina Corradi, Eduardo Mendez-Villuendas, Helgi I Ingólfsson, Ruo-Xu Gu, Iwona Siuda, Manuel N Melo, Anastassiia Moussatova, Lucien J DeGagné, Besian I Sejdiu, Gurpreet Singh, et al. Lipid-protein interactions are unique fingerprints for membrane proteins. ACS Central Science, 4(6):709–717, 2018.

[43] Besian I Sejdiu and D Peter Tieleman. Lipid-protein interactions are a unique property and defining feature of G Protein-Coupled receptors. Biophysical Journal, 118(8):1887–1900, 2020.

[44] William Humphrey, Andrew Dalke, and Klaus Schulten. VMD: visual molecular dynamics. Journal of Molecular Graphics, 14(1):33–38, 1996.

[45] Mikhail A. Lomize, Andrei L. Lomize, Lrina D. Pogozheva, and Henry I. Mosberg. OPM: Orientations of proteins in membranes database. Bioinformatics, 22:623–625, 2006.

[46] Tsjerk A. Wassenaar, Helgi I. Ingólfsson, Rainer A. Böckmann, D. Peter Tieleman, and Siewert J. Marrink. Computational lipidomics with insane: A versatile tool for generating custom membranes for molecular simulations. Journal of Chemical Theory and Computation, 11:2144–2155, 2015.

[47] Berk Hess, Carsten Kutzner, David van der Spoel, and Erik Lindahl. Gromacs 4: Algorithms for highly efficient, load-balanced, and scalable molecular simulation. Journal of Chemical Theory and Computation, 4:435–447, 2008.

[48] David van der Spoel, Erik Lindahl, Berk Hess, Gerrit Groenhof, Alan E. Mark, and Herman J. C. Berendsen. Gromacs: Fast, flexible, and free. Journal of Computational Chemistry, 26:1701–1718, 2005.

[49] Djurre H. De Jong, Gurpreet Singh, W. F. Drew Bennett, Clement Arnarez, Tsjerk A. Wassenaar, Lars V. Schäfer, Xavier Periole, D. Peter Tieleman, and Siewert J. Marrink. Improved parameters for the martini coarse-grained protein force field. Journal of Chemical Theory and Computation, 9(1):687–697, 2013.

[50] Giovanni Bussi, Davide Donadio, and Michele Parrinello. Canonical sampling through velocity rescaling. Journal of Chemical Physics, 126(1):014101, 2007.

[51] H. J. C. Berendsen, J. P. M. Postma, W. F. van Gunsteren, A. DiNola, and J. R. Haak. Molecular dynamics with coupling to an external bath. Journal of Chemical Physics, 81(8):3684–3690, 1984.

[52] W. Helfrich. Elastic properties of lipid bilayers: theory and possible experiments. Z. Naturforsch., 28:693–703, 1973.

[53] Frank LH Brown. Elastic modeling of biomembranes and lipid bilayers. Annual Review of Physical Chemistry, 59:685–712, 2008.

[54] Erik Lindahl and Olle Edholm. Mesoscopic undulations and thickness fluctuations in lipid bilayers from molecular dynamics simulations. Biophysical Journal, 79(1):426–433, 2000.

[55] Philip W. Fowler, Jean Hélie, Anna Duncan, Matthieu Chavent, Heidi Koldsø, and Mark S. P. Sansom. Membrane stiffness is modified by integral membrane proteins. Soft Matter, 12(37):7792–7803, 2016.

[56] Antreas C Kalli and Reinhart AF Reithmeier. Interaction of the human erythrocyte Band 3 anion exchanger 1 (ae1, slc4a1) with lipids and glycophorin A: molecular organization of the Wright (wr) blood group antigen. PLoS Computational Biology, 14(7):e1006284, 2018.

[57] Diomedes E Logothetis, Vasileios I Petrou, Miao Zhang, Rahul Mahajan, Xuan-Yu Meng, Scott K Adney, Meng Cui, and Lia Baki. Phosphoinositide control of membrane protein function: a frontier led by studies on ion channels. Annual Review of Physiology, 77:81–104, 2015.

[58] Camilla Raiborg, Eva M Wenzel, Nina M Pedersen, and Harald Stenmark. Phosphoinositides in membrane contact sites. Biochemical Society Transactions, 44(2):425–430, 2016.

[59] Peter J Hamilton, Andrea N Belovich, George Khelashvili, Christine Saunders, Kevin Erreger, Jonathan A Javitch, Harald H Sitte, Harel Weinstein, Heinrich JG Matthies, and Aurelio Galli. PIP 2 regulates psychostimulant behaviors through its interaction with a membrane protein. Nature Chemical Biology, 10(7):582–589, 2014.

[60] George Hedger, David Shorthouse, Heidi Koldso, and Mark S. P. Sansom. Free energy landscape of lipid interactions with regulatory binding sites on the transmembrane domain of the EGF receptor. Journal of Physical Chemistry B, 120:8154–8163, 2016.

