## Supplementary Figures and Tables for "Lipid-mediated organization of prestin in the outer hair cell membrane and its implications in sound amplification"

### Supporting Information

Table S1: ***Quad-prestin* simulations.** Each system included four copies of prestin dimer arranged in a square configuration with a side length of 100 Å and simulated for 20 $\mu$s, resulting in a collective sampling time of 80 $\mu$s (4 dimers $\times$ 20 $\mu$s = 80 $\mu$s) of lipid-protein interactions per prestin conformation (contracted or expanded) and an overall simulation time of 160 $\mu$s.

| System | Conformation | Number of dimers | Simulation time per dimer | Total sampling time |
|---|---|---|---|---|
| 1 | Contracted | 4 | 20 $\mu$s | 80 $\mu$s |
| 2 | Expanded | 4 | 20 $\mu$s | 80 $\mu$s |

Table S2: ***Double-prestin* simulations.** Each system included a pair of prestin dimers, either in contracted on in expanded conformations, in one of the 16 different relative orientations between the two dimers, and simulated for $10\,\mu$s, resulting collectively in $320\,\mu$s (16 orientations × 2 conformations × $10\,\mu s = 320\,\mu s$) of simulation time.

| System | Conformation | Orientation of Dimer I | Orientation of Dimer II | Simulation time |
|---|---|---|---|---|
| 1 | Contracted | 0 | 0 | $10\,\mu$s |
| 2 | Contracted | 0 | 45 | $10\,\mu$s |
| 3 | Contracted | 0 | 90 | $10\,\mu$s |
| 4 | Contracted | 0 | 135 | $10\,\mu$s |
| 5 | Contracted | 45 | 0 | $10\,\mu$s |
| 6 | Contracted | 45 | 45 | $10\,\mu$s |
| 7 | Contracted | 45 | 90 | $10\,\mu$s |
| 8 | Contracted | 45 | 135 | $10\,\mu$s |
| 9 | Contracted | 90 | 0 | $10\,\mu$s |
| 10 | Contracted | 90 | 45 | $10\,\mu$s |
| 11 | Contracted | 90 | 90 | $10\,\mu$s |
| 12 | Contracted | 90 | 135 | $10\,\mu$s |
| 13 | Contracted | 135 | 0 | $10\,\mu$s |
| 14 | Contracted | 135 | 45 | $10\,\mu$s |
| 15 | Contracted | 135 | 90 | $10\,\mu$s |
| 16 | Contracted | 135 | 135 | $10\,\mu$s |
| 17 | Expanded | 0 | 0 | $10\,\mu$s |
| 18 | Expanded | 0 | 45 | $10\,\mu$s |
| 19 | Expanded | 0 | 90 | $10\,\mu$s |
| 20 | Expanded | 0 | 135 | $10\,\mu$s |
| 21 | Expanded | 45 | 0 | $10\,\mu$s |
| 22 | Expanded | 45 | 45 | $10\,\mu$s |
| 23 | Expanded | 45 | 90 | $10\,\mu$s |
| 24 | Expanded | 45 | 135 | $10\,\mu$s |
| 25 | Expanded | 90 | 0 | $10\,\mu$s |
| 26 | Expanded | 90 | 45 | $10\,\mu$s |
| 27 | Expanded | 90 | 90 | $10\,\mu$s |
| 28 | Expanded | 90 | 135 | $10\,\mu$s |
| 29 | Expanded | 135 | 0 | $10\,\mu$s |
| 30 | Expanded | 135 | 45 | $10\,\mu$s |
| 31 | Expanded | 135 | 90 | $10\,\mu$s |
| 32 | Expanded | 135 | 135 | $10\,\mu$s |

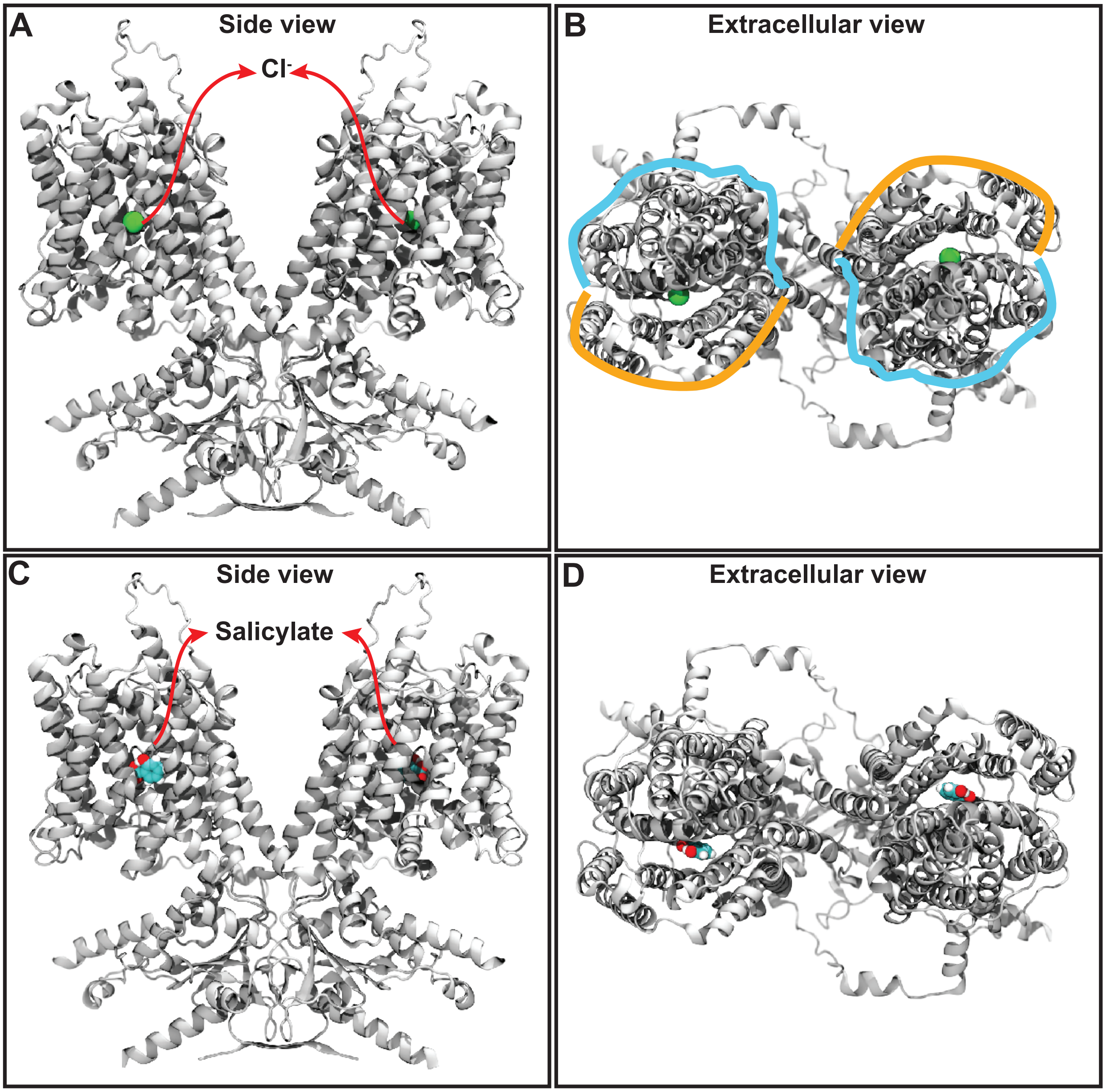


Figure S1: **All-atom models of prestin in different functional (conformational) states.** **(A/B)** Membrane and extracellular views of contracted prestin bound to $Cl^-$ (green). The core and gate domains are highlighted in blue and orange, respectively. **(C/D)** Membrane and extracellular views of expanded prestin bound to the inhibitor salicylate. Both ligands, $Cl^-$ and salicylate, bind to the same site, located in between the core and gate domains. The contracted conformation has a smaller cross-sectional area in the membrane.

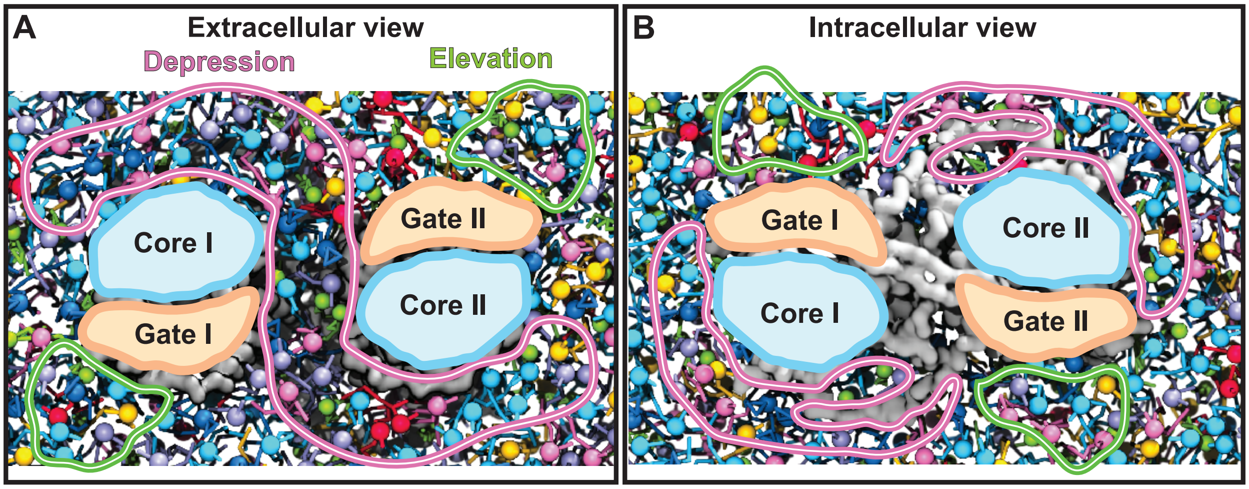


Figure S2: **Membrane deformation pattern around prestin.** Extracellular **(A)** and intracellular **(B)** views of a membrane-embedded prestin dimer after 20 $\mu$s of simulation. Regions with significant lipid elevation (green) or depression (pink) are highlighted in the vicinity of the gate (orange) and core (blue) domains, respectively. PC, PI, SM, CHOL, PS, PG, and PE lipids are shown in cyan, blue, magenta, green, purple, red, and yellow, respectively.

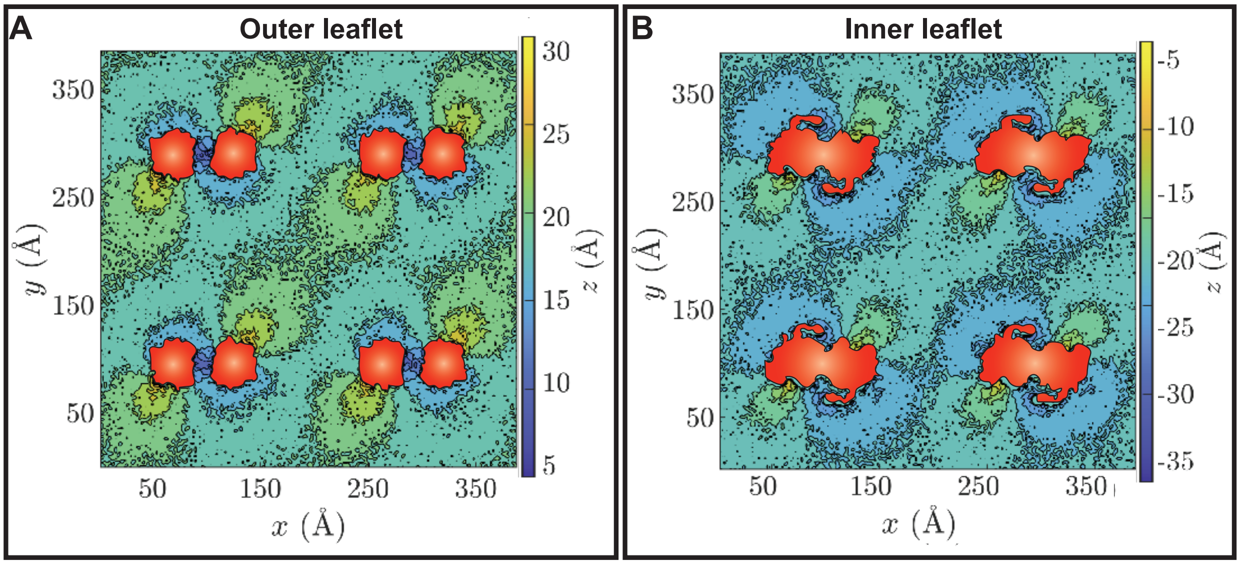


Figure S3: **Prestin-induced, anisotropic membrane deformation for the expanded conformation.** 2D histograms of the $z$ positions of lipid headgroups for the outer **(A)** and inner **(B)** leaflets. Both leaflets are viewed from the extracellular side. Histograms are constructed from the last 5 $\mu$s of a 20-$\mu$s trajectory. The cross sectional areas of the proteins in each leaflet are drawn in red.

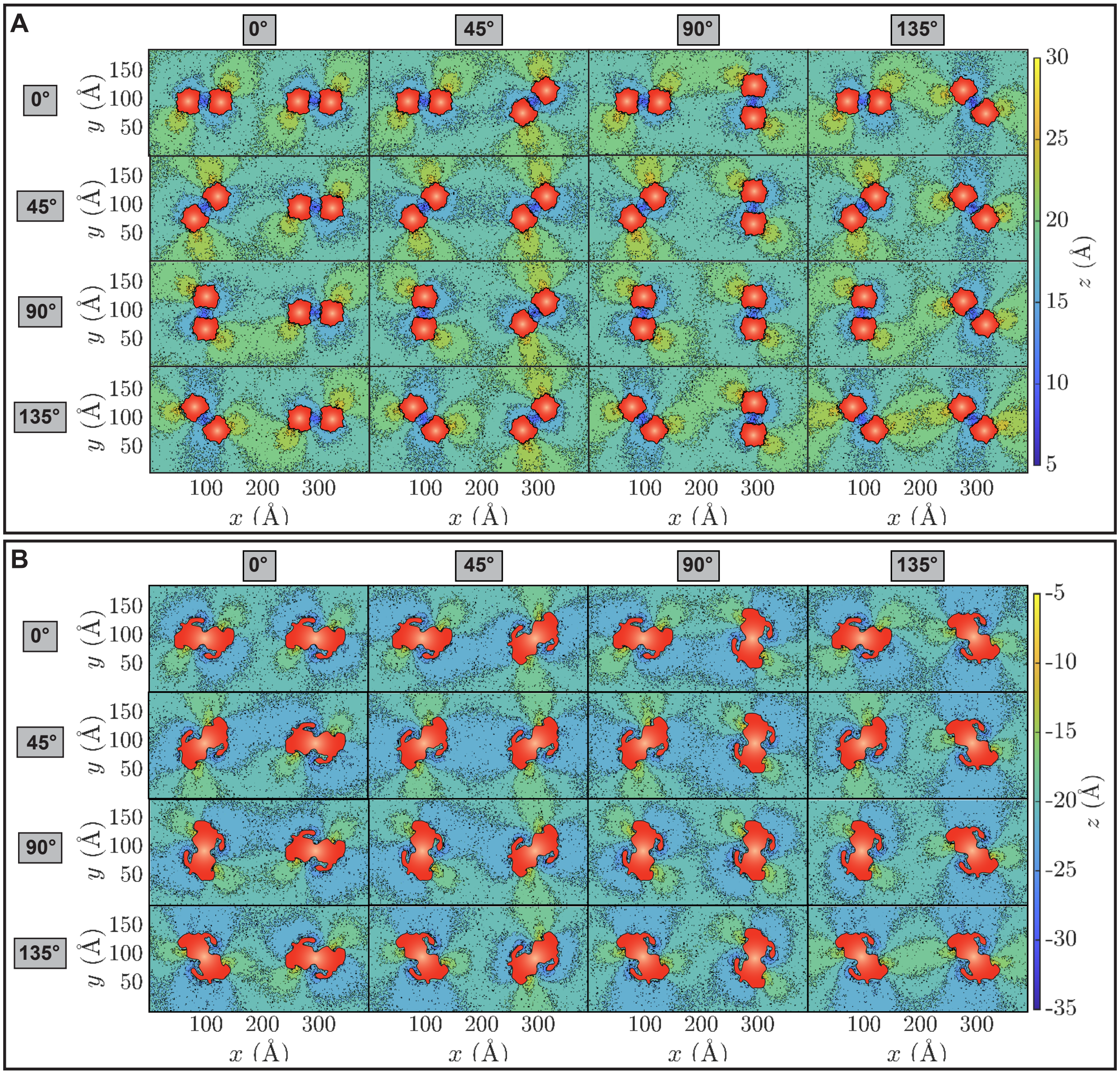


Figure S4: **Constructive or destructive interference of deformation patterns induced by expanded prestin's different orientations**. Heatmaps of phospholipid height in the outer **(A)** and inner **(B)** leaflets are calculated using the procedure described in Fig. 2. The prestin dimers are placed at $(x, y) = (100, 100)$ Å and $(300, 100)$ Å, respectively. The orientation of Dimer I (angle with respect to the $x$ axis) is specified on the left, and for Dimer II on the top of the panel. The protein cross sectional area in each leaflet is drawn in red.

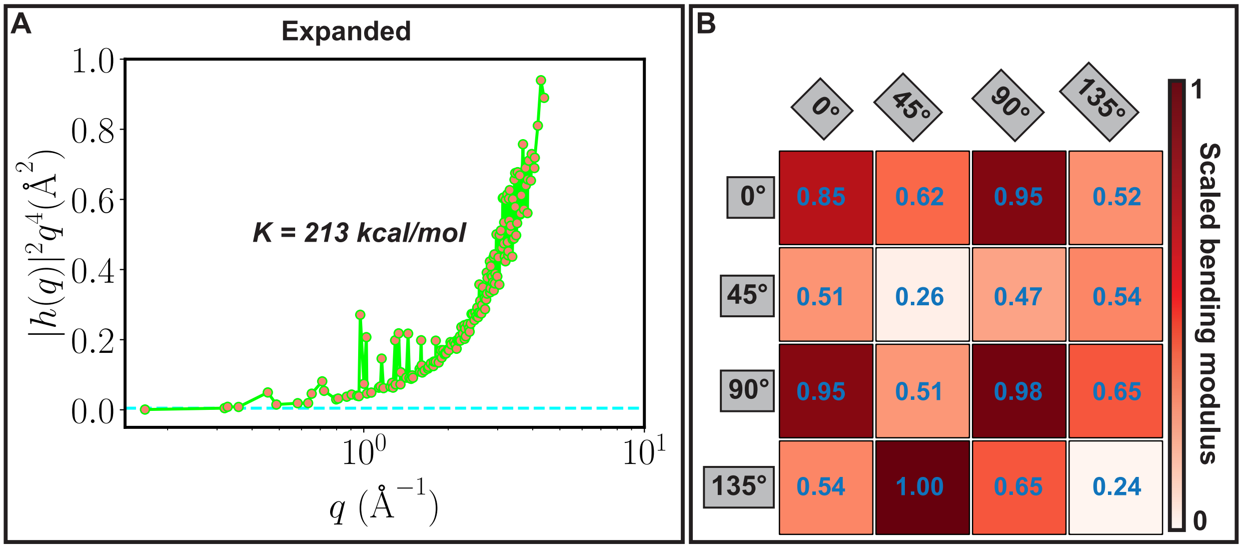


Figure S5: **Membrane bending moduli calculated for different expanded prestin configurations**. **(A)** $|h(q)|^2q^4$ plotted vs. $q$, wavenumber, for the system with two prestin dimers both at 0° ([0°, 0°]). The $y$ value of the horizontal line fitted to the function at low $q$ values (blue dashed line) is equivalent to $\frac{k_B T}{K}$, and is used to calculate $K$. For the represented system here, $K = 213\,\text{kcal/mol}$. **(B)** Bending moduli for the membranes with two prestin dimers in different orientations (Fig. 3), normalized by the maximum value for the [45°, 135°] system. The orientation of Dimer 1, placed at $(x, y) = (100, 100)$ Å is shown on the left, and for Dimer II, placed at $(x, y) = (300, 100)$ Å, on the top of the table. The color shade in each box also represents the strength of the bending modulus, and shown in the scale bar. The minimum bending moduli are captured for the system with two prestin dimers at either 45° or 135°.

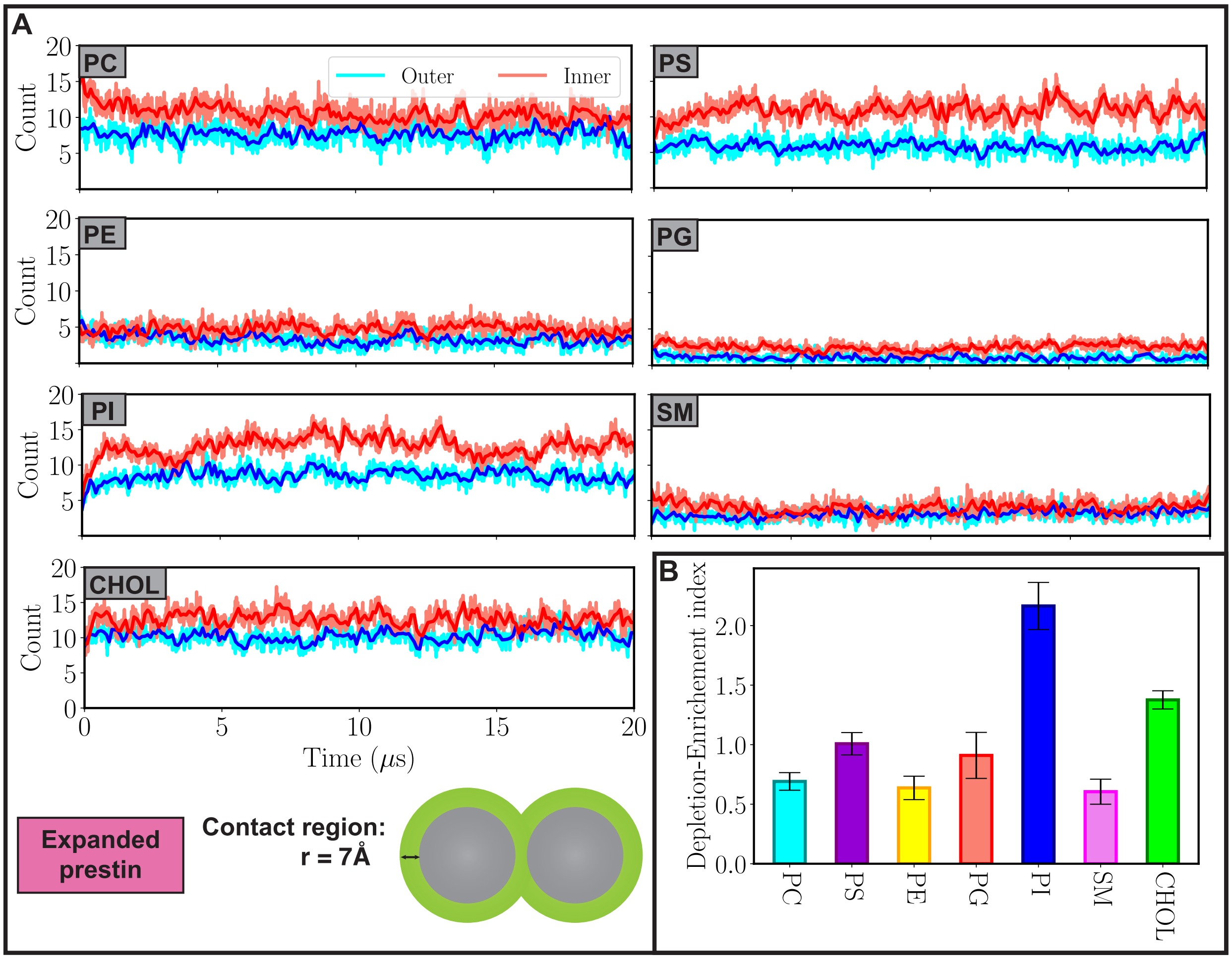


Figure S6: **Enrichment/depletion of different lipids around expanded prestin**. **(A)** Time series of the number of different lipid types within 7 Å of prestin, averaged over four dimers, in the outer (blue) and inner (red) leaflets. PI lipids accumulate the most around prestin in the inner leaflet. **(B)** Depletion-Enrichment index of each lipid type (the ratio of the local and bulk fractions of the lipid). PI and CHOL show the highest enrichment, whereas SM and PE show largest extents of depletion.
